# How did spotted hyaenas respond to decreased prey availability in their clan territories over the last decades?

**DOI:** 10.64898/2026.02.10.705143

**Authors:** Mellina Sidous, Morgane Gicquel, Sonja Metzger, Marion L. East, Heribert Hofer, Julius W. Nyahongo, Sarah Benhaiem, Sarah Cubaynes

## Abstract

Climate change can alter predator-prey dynamics by influencing the distribution and movements of migratory prey. Despite increasing research on predator-prey mismatches, how predators respond to changes in prey availability caused by climate change remains largely unknown, particularly for behaviourally flexible species such as central-place foragers. In the Serengeti National Park, Tanzania, increased rainfall in recent decades is thought to have altered the movement patterns of large herds of migratory herbivores, the main prey of spotted hyaenas (*Crocuta crocuta*) in the park, leading to a decrease in yearly migratory prey presence within hyaena clan territories. Between 1990 and 1994, migratory prey were present within hyena clan territories for about 12% of the year—mainly in May–June and November-December—dropping to just 7% between 2015 and 2019. Using longitudinal data from three Serengeti hyaena clans at the centre of the park monitored between 1990 and 2019, we investigated the impact of the observed decrease in migratory prey presence on the number of observations of hyaenas feeding at carcasses (“feeding events”, n = 777), and explored associated changes in hyaena clan size. The vast majority of observed hyaena feeding events involved migratory prey species, with this relative proportion remaining constant throughout the study period. Annual patterns in the number of feeding events closely mirrored annual patterns of migratory prey presence in clan territories, with two distinct peaks mid-year and toward the end of the year. As migratory prey presence in the study clan territories declined over the years, the number of observed feeding events also decreased. However, the size of two out of three clans increased over time, suggesting that the decline in migratory prey presence in clan territories and in the number of observed feeding events did not negatively impact hyaena clans. This absence of decline in clan size may reflect the fact that hyaenas feed within their territories for only a small fraction of the year, although it also invites further investigation into the mechanisms hyaenas may employ to compensate for reduced prey availability and reduced feeding events within their clan territories.

## Introduction

Climate-change driven alterations in rainfall or temperature can affect the phenology, spatial distribution, behaviour, and demography of animal populations (Lecomte et al. 2009, Hetem et al. 2014, Iler et al. 2021). These changes can cascade through ecological communities, and modify the dynamics of interspecific interactions, such as predator-prey relationships. Climate-driven resource limitations can compel prey species to shift habitat use (Tablado et al. 2014) or activity level (Anholt and Werner 1998, Lecomte et al. 2009), and may affect prey body condition, survival, and ultimately abundance (Labadie et al. 2023). Such changes in prey populations can, in turn, affect predator-prey encounter rates and the hunting success of predators (Lecomte et al. 2009, Morin et al. 2021), and may alter the frequency of predation events. If predators are resource-limited, a decline in predation events frequency can decrease individual predator body condition (Stewart et al. 2021) and trigger population decline either through demographic processes (Schmidt et al. 2012), or space use mechanisms, such as an increase in home ranges (Mills and Knowlton 1991, Schmidt 2008).

In addition to numerical responses, predators may adjust their behaviour to cope with reduced predation opportunities by shifting their diet to different prey (Cooper et al. 1999, Elbroch et al. 2013, Labadie et al. 2023), increasing scavenging (Höner et al. 2002, Roth 2003), or by increasing activity and foraging effort (Ronconi and Burger 2008, Schmidt 2008). Although these behaviours may buffer the negative effects of reduced predator-prey encounter rates and hunting success, numerical declines in predator populations may still occur (Ronconi and Burger 2008, Schmidt 2008). Even though various behavioural and demographic responses to changing prey availability have been documented, the dynamics of such responses remain poorly understood in long-distance central-place foragers, such as sea lions (*Arctocephalus gazella;* Jeanniard-du-Dot et al. 2017), wolves (*Canis lupus*; Walton et al. 2001) or spotted hyaenas (*Crocuta crocuta;* Hofer and East 1993a). Such predators must commute between a fixed site and distant, shifting foraging areas, due to the mobility of their main prey. Here, we explore how a climate change-driven reduction in prey availability in the territories of three clans of spotted hyaenas (hereafter referred to as hyaenas) in the Serengeti National Park, Tanzania, (Gicquel et al. 2022b) has affected the number of feeding events, i.e. the observations of hyaenas feeding at carcasses within their clan territories and clan size.

The Serengeti National Park, covering approximately 14,750 km², is part of the larger Serengeti–Mara ecosystem, which spans northern Tanzania and southwestern Kenya. This ecosystem is defined by the seasonal migration of three herbivore species, blue wildebeest (*Connochaetes taurinus*), plains zebra (*Equus quagga*) and Thomson’s gazelle (*Gazella thomsoni*), which track seasonal shifts in grazing resource availability that are shaped by spatial and temporal rainfall patterns across the region (Wilmshurst et al. 1999, Boone et al. 2006). During the wet season, migratory prey concentrates in the open, nutrient-rich short-grass plains of the south-eastern Serengeti to calve. As the dry season begins around mid-May to early June and the southern plains dry out, herbivores gradually move northwest toward the Mara and north-western Serengeti’s wooded grasslands, where annual rainfall is about twice as high as that of southern plains and vegetation biomass remains higher (Boone et al. 2006). Migratory prey remain in north-western Serengeti until around October, before returning south with the onset of the rains (see Figure 1, Panel A).

**Figure 1:**
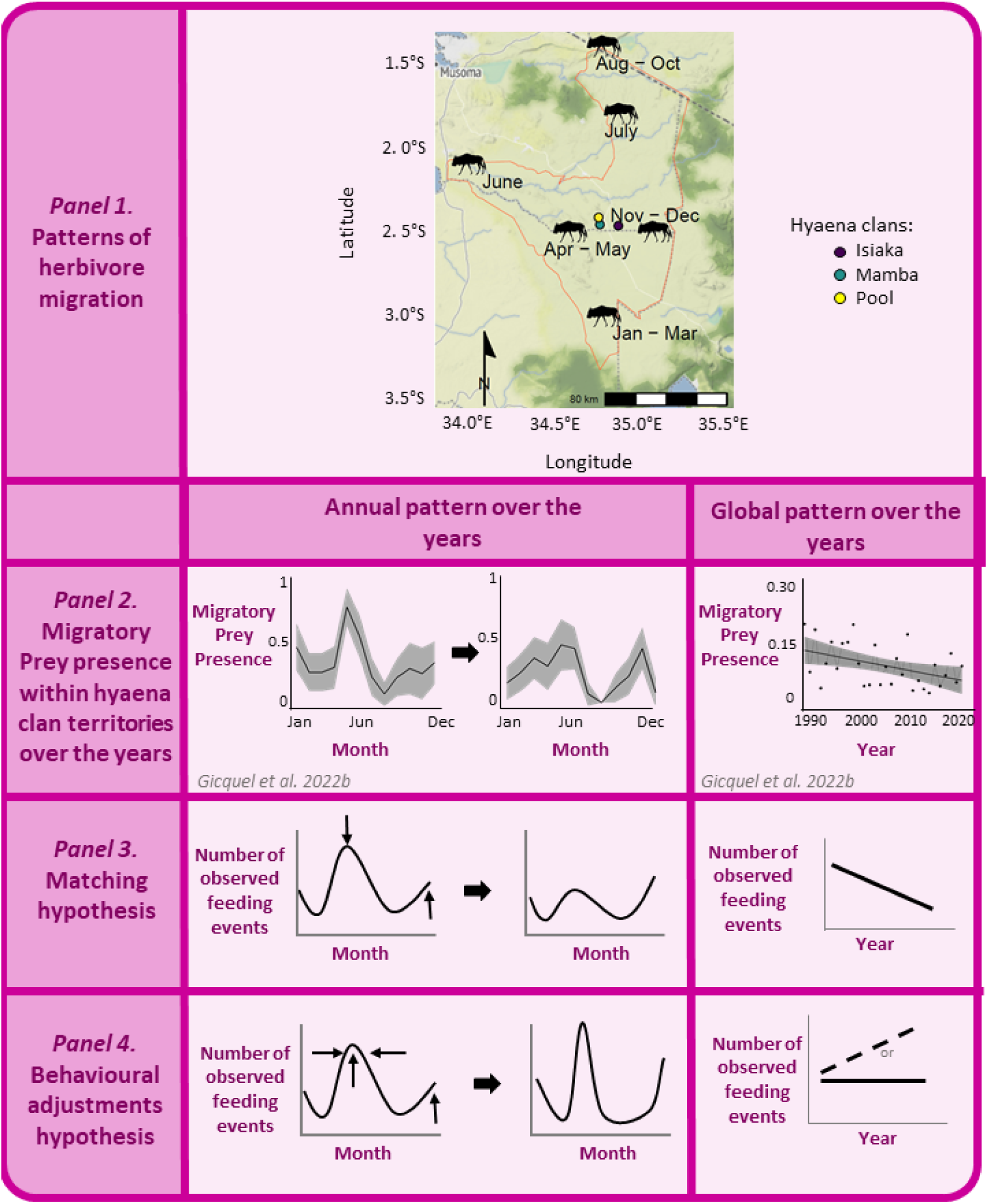
Patterns of migratory prey presence in hyaena clan territories and main hypotheses. **Panel 1:** Migration route of migratory prey (wildebeest silhouettes) in the Serengeti National Park throughout the year, and location of the three hyaena clan studied. Silhouette of *Connochaetes taurinus* from S. Metzger. **Panel 2.** Patterns of migratory prey presence in the territories of the three studied hyaena clans across decades. Figure from Gicquel et al. (2022b). **Panel 3:** Matching hypothesis, i.e. a temporal association between migratory prey presence and the number of observed hyaena feeding events. Patterns of the number of observed feeding events are expected to mirror those of migratory prey presence, with a reduction in the May-June peak, leading to fewer feeding events observed over the years. **Panel 3:** Compensation through behavioural adjustments hypothesis. Over the years, we expect hyaenas to have increased their foraging effort during periods of high migratory prey presence in their clan territories, namely in May-June and November-December, to compensate for the overall reduced migratory prey presence. If the number of observed feeding events increased in those months, we predict a stable or slightly increasing number of feeding events over the years, despite the declining migratory prey presence

In the Serengeti, migratory herbivores dominate the large mammalian herbivore community, and hyaenas at the centre of the park depend on them, as resident prey populations in this area are insufficient to feed the entire clans (Hofer and East 1993a). To cope with the large fluctuations in prey availability caused by prey migration, hyaenas employ a distinctive foraging strategy. The long-term monitoring of three hyaena clans located in central Serengeti showed that clans defend a fixed territory, where all members feed when migratory prey are present and locally abundant (∼238.5 animals/km²), typically in May–June and November–December (see Figure 1, Panel 2). However, periods of high migratory prey presence in clan territories occur only 2-20% of the year (Appendix S1). For the remainder of the year, when resident prey density drops down to ∼7.2 animals/km², all clan members regularly undertake “commuting trips”—foraging excursions that can cover up to 70 km and last several days—to reach areas with greater prey availability, i.e. where migratory prey are located (Hofer and East 1993a, b; Hofer et al. 1995). Although commuting is also observed in other hyaena populations (e.g. Höner et al. 2005, Holekamp and Dloniak 2011), hyaenas in the Serengeti are one of the most outstanding example of behavioural flexibility and adjustment to frequent and large fluctuations in prey availability, as they undertake unusually long and distant commuting trips (Hofer and East 1993a, 1993b, 1993c, Hofer et al. 1995).

In recent decades, increasing evidence has suggested a shift in the movement patterns of migratory prey in the Serengeti, linked to a documented increase in both wet- and dry-season rainfall over the past 30 years (Mahony 2020, Gicquel et al. 2022b, Mtewele et al. 2023, Ogutu et al. 2024). For the three hyaena clans monitored in central Serengeti, migratory prey presence within territories declined from a mean of 12% of the time in the first five years of monitoring to 7% of the time in the last five years (see Appendix S1). This decline was most pronounced in May–June, a period historically supporting high densities of migratory prey within clan territories with all clan members typically feeding locally, i.e. at kills in their clan territories (Gicquel et al. 2022a; Figure 1, Panel 2). This decline suggests that climate change is challenging within-territory feeding opportunities for the studied clans.

Given the decline in migratory prey presence within clan territories during the last three decades, one might expect hyaenas to compensate by increasing their commuting effort. However, the presence of lactating females at communal dens remained constant over the last 30 years, suggesting no changes in commuting effort and that hyaenas may not have left their clan territories more often or for longer periods (Gicquel et al. 2022b). This finding raises the question: how did hyaenas respond to the overall decline in migratory prey presence within their clan territories? Did the number of feeding events decline alongside migratory prey presence, and how did clan size vary over the last three decades? To investigate these questions, we investigated temporal patterns of incidental observations of hyaena feeding events recorded over 30 years, arising either from predation or scavenging, and recorded in the territories of the three studied clans. We also took advantage of the long-term monitoring of the size of these clans to explore temporal trends and in particular ask whether the reduction in migratory prey presence could be associated with demographic costs.

We propose two hypotheses to explain patterns in the number of feeding events and clan size (Figure 1). First, we could expect a positive association between migratory prey presence and the number of observed hyaena feeding events within clan territories. In this “matching hypothesis”, the number of feeding events is driven by the presence of migratory prey within hyaena clan territories (Hofer and East 1993a). Accordingly, feeding events patterns should mirror the patterns of migratory prey presence in hyaena clan territories reported by Gicquel et al. (2022a) and shown in Figure 1, Panel 2. Specifically, we expect to observe: i) a change in the phenology of feeding events on an annual scale, with a reduced mid-year (May-June) peak; and ii) fewer feeding events over the years (Figure 1, Panel 3). Alternatively, the “compensation through behavioural adjustments” hypothesis proposes that hyaenas attempt to compensate for the reduced migratory prey presence over the years by increasing foraging effort during peak months (May-June and November-December), i.e. by killing more individual migratory prey, to take advantage of relatively easily available prey. Under this hypothesis, the number of feeding events in these peak months should increase over time (Figure 1, Panel 4). If this increased foraging effort during peak periods offsets the overall decline in migratory prey presence, we might observe a stable or even slightly increasing number of feeding events over the years.

We also examine trends in clan size over the study period and explore a potential positive association with the number of feeding events. Since lactating females do not appear to increase commuting effort in response to reduced migratory prey availability (Gicquel et al. 2022b), reduced within-territory feeding could reflect a globally slightly reduced food intake, potentially affecting milk transfer and offspring body condition, survival and ultimately clan size. However, because hyaenas feed within their own clan territories only during limited periods of the year, they are not solely dependent on feeding events within territories. Access to prey outside of territories during the many months when migratory herds are absent plays a critical role to determine cub growth and survival and therefore hyaena clan size. Thus, although worth investigating, we did not strongly expect to observe a positive association between within-territory feeding events and clan size. Nonetheless, we only disposed of feeding observations within clan territories, offering a valuable snapshot of one key area where feeding occurs and provide insight into how hyaenas may be responding to local changes in resource availability. Understanding within-territories feeding patterns provides part of the picture of the broader foraging strategy of hyaenas, and brings insight into the extent to which local changes in prey availability might affect clan size in a long-distance central-place forager.

## Material and Methods

### Data collection and definition of variables

The Serengeti ecosystem straddles the border between Tanzania and Kenya, directly east of Lake Victoria in East Africa. Within this ecosystem, the Serengeti National Park is located in northwestern Tanzania, East Africa. Data were collected in the context of a long-term research project focused on three spotted hyaena clans that held permanent territories in the centre of the park, halfway between the dry- and wet-season ranges of migratory prey (2° 26’ S, 34° 47’ E; Hofer and East 1993b) that have been monitored for more than 35 years (Isiaka [I] since May 1987, Pool [P] since October 1989, and Mamba [M] since August 1990). Communal den(s)—the social centres of hyaena clans (Frank 1986a, b, Strauss et al. 2024) — are regularly visited to record the presence of clan members within a 100 m radius and social interactions. Individuals were recognized based on distinct spot patterns, scars and ear notches (Frank 1986b, Hofer and East 1993a). Clans comprise immigrant and natal breeding males, multiple generations of philopatric females, and their dependent offspring (Frank 1986b). Offspring, observable shortly after birth while being nursed at underground burrow entrances, are sexed by the age of three months based on morphological differences in the glans of the erect phallus (Frank et al. 1991). Each clan was monitored for approximately 2-3h at dawn and dusk, when the presence of individuals at communal dens was highest (Hofer and East 1993a, 1993b, 1993c, Gicquel et al. 2022b).

We focused on monitoring sessions conducted from January 1990 to December 2019. To ensure sufficient data while maintaining a relatively fine-scale temporal resolution, data were aggregated monthly by summing values, computing means or medians, or using the maximum or most frequently recorded value, as specified for each variable below.

#### Observation effort

Over the 30-year study period (January 1990 to December 2019), 360 months elapsed. All clans were monitored at least one day per month during more than 80% of these months (329, 302 and 321 months for Isiaka, Mamba, and Pool, respectively). On average, each clan was monitored 11.32 days per month (standard deviation σ = 28.31), without differentiating between days with one (morning or evening) or two (morning and evening) monitoring sessions. The maximum number of monitored days per month was 28 in Isiaka, 29 in Pool, and 24 in Mamba. Since the number of days monitored varied from month to month across clans, we computed the variable *n_monitored_*, the number of monitoring sessions (i.e., days monitored) per clan per month, to account for variation in observation effort in the analyses.

#### Incidental observations of hyaena feeding at carcasses (feeding events)

During each monitoring session, observers recorded any incidental sighting of hyaena feeding events within the territory of the monitored clan. We defined feeding events as any occasion during which hyaenas from one of the three study clans were observed feeding on a carcass, whether they killed the prey or scavenged it and regardless of the species, sex and age of prey. In addition, observers recorded feeding events outside of monitoring sessions, on the way to a given clan territory, or on the way back to the research station. when encountered during routine observations. For each observation, observers recorded the date, GPS location, hyaena clan territory, species, age, sex and cause of prey death if possible, presence of other carnivores, and the number of hyaenas present and their identities. A total of 777 feeding events were observed.

Based on this dataset, we computed the total number of incidental feeding events recorded each month of the study period in each clan territory (*n_feeding;_* ; see Appendix S2, Table 1 for list of variables abbreviations). The number of feeding events observed each month in each clan (*n_feeding_*) ranged from 0 to 10 and was 0.79 (σ = 1.62) on average. When a clan was visited at least once during a month but no feeding event was observed, a value of 0 was assigned to *n_feeding_* to differentiate this case from month(s) when the clan was not visited at all, for which a value NA was attributed. We identified the clan in which each feeding event was seen in a categorical variable *Clan* (levels: I for Isiaka, M for Mamba, P for Pool).

For 732 observed feeding events, we could determine whether the prey consumed was a migratory species (blue wildebeest, Thomson’s gazelle, or plain zebra; see Appendix S3 for the list of resident species), recorded in a variable called *“Prey_type”* (levels: 0 = resident prey; 1 = migratory prey). Migratory prey accounted for 82% of all observations (*n* = 732). See Appendix S3 for further details on *Prey_type*, and differences between clans.

#### Migratory prey presence

Migratory prey presence within hyaena clan territories (*Prey*) was inferred using a categorical index of prey abundance previously defined for this area, based on transect data (Hofer and East 1993a). It took a value of 1 if migratory prey were absent (∼7.2 (resident) prey individuals per km^2^), 2 if present in low numbers (∼31 prey individuals per km^2^), and 3 if abundant (∼238.5 prey individuals per km^2^_;_ see methods in Hofer and East 1993a, b, a).

We summarize monthly trends in migratory prey presence, using two additional indices: the most common (*Prey_mode_*) and the maximum (*Prey_max_*) value of *Prey* for a given month and clan. We explored *Prey_max_* based on the rationale that if migratory prey were observed as abundant in a clan’s territory, they may have been present for a few days within or near its boundaries. Hence, a *Prey_max_* score of 3 possibly indicates that prey was overall close to hyaena clan territories during this month, and thus, that they had to travel little to feed.

#### Clan size

During each monitoring session, the presence of all clan members was recorded, which, along with life history data (birth, emigration and death dates), enabled the estimation of clan size (number of alive members of a given clan) and composition throughout the study period (e.g., Gicquel et al. 2022b, a, Naciri et al. 2023). Clan size was defined here as the median number of alive clan members per clan each month. Due to the high frequency of den observations and reliable individual identification, uncertainty in population size estimates was minimal.

### Statistical analyses

We used General Linear Models (GLMs; for linear, categorical, or quadratic effects; package *MASS*, v7.3 ; Venables et al. 2002) and Generalized Additive Models (GAMs; for nonlinear effects; package *gam*, v1.2; Hastie 2004) using software R version 4.2.2 (R Core Team 2022) to explore the predictors of *n_feeding_* and *Prey_type* in hyaena clan territories. We performed overdispersion tests using the *DHARMa* package (v0.4.6; Hartig 2016), which indicated that a negative binomial distribution was more suitable than a Poisson distribution for modelling *n_feeding_* (see Appendix S4). Models exploring the predictors of *Prey_type* were run using a binomial distribution. Statistical analyses of *Prey_type* confirmed that hyaenas could not compensate for the decline in migratory prey presence in their territory over the years by shifting to alternative prey types, as the proportion of migratory prey among observed feeding events showed no decrease or trend over time (Appendix S5).

We used a model selection approach based on the Akaike Information Criterion (AIC; Burnham and Anderson 2010), with two separate selections for models predicting *n_feeding_* and *Prey_type*. An effect of *Clan* was tested in all models unless stated otherwise. We considered two models equivalent when ΔAIC <5 (Burnham and Anderson 2010). We also reported the percentage of deviance explained by each model (R^2^ in Appendix S5, Table S1 and Appendix S6, Table S1). We computed estimates using the function ggpredict from the *ggeffects* package (v1.5.2; Lüdecke 2018) with 95% confidence intervals (CIs) shown in brackets. Figures were plotted using the *ggplot2* package (v3.43; Wickham 2016).

We first investigated monthly variation (also termed annual patterns) in *n_feeding_* averaged over the entire study period using quadratic (*qMonth*), categorical (*cMonth*), or nonlinear (*sMonth*) effects of the variable *Month* as a predictor variable. We then included an effect of *Year* to investigate how *n_feeding_* varied over the past decades. We tested linear (*lYear*), quadratic (*qYear*), categorical (*cYear*), random (*rYear*), and nonlinear (*sYear*) effects of *Year*, and tested whether annual patterns remained unchanged or varied over time by including models with an interaction between the effects of *Year* and *Month*.

We also added *Prey* (*Prey_max_* and *Prey_mode_*) as categorical explanatory variables to test whether the index of migratory prey presence in hyaena territories was a good predictor of *n_feeding_*. We included *n_monitored_* as an offset in all models exploring the predictors of *n_feeding_* to account for potential variation in observation effort.

## Results

### Temporal patterns in the number of feeding events observed

#### Annual pattern: two main peaks in observed feeding events

We identified a clear seasonal pattern in the number of observed feeding events, with lower numbers of observed feeding events during the first months of the year (between January and June) than later months (Figure 2A; see Appendix S7 for results without a *Clan* effect). As expected under both the matching and behavioural adjustment hypotheses (Figure 1, Panel 3 and 4), the number of observed feeding events peaked slightly around mid-year (May-June) and toward the end of the year (November-December) — the two periods during which migratory prey are typically abundant within hyaena clan territories (estimates from *cMonth* and *sMonth* models; ΔAIC = 0 and 0.05 among models exploring annual patterns; Appendix S6). All clans exhibited similar annual patterns, although Pool showed consistently higher values during the second part of the year, from June to December—34% and 42% higher than Isiaka and Mamba, respectively, in December (Figure 2A; estimates from the *sMonth* * *Clan* model; ΔAIC = 4.8 among models exploring annual patterns).

**Figure 2:**
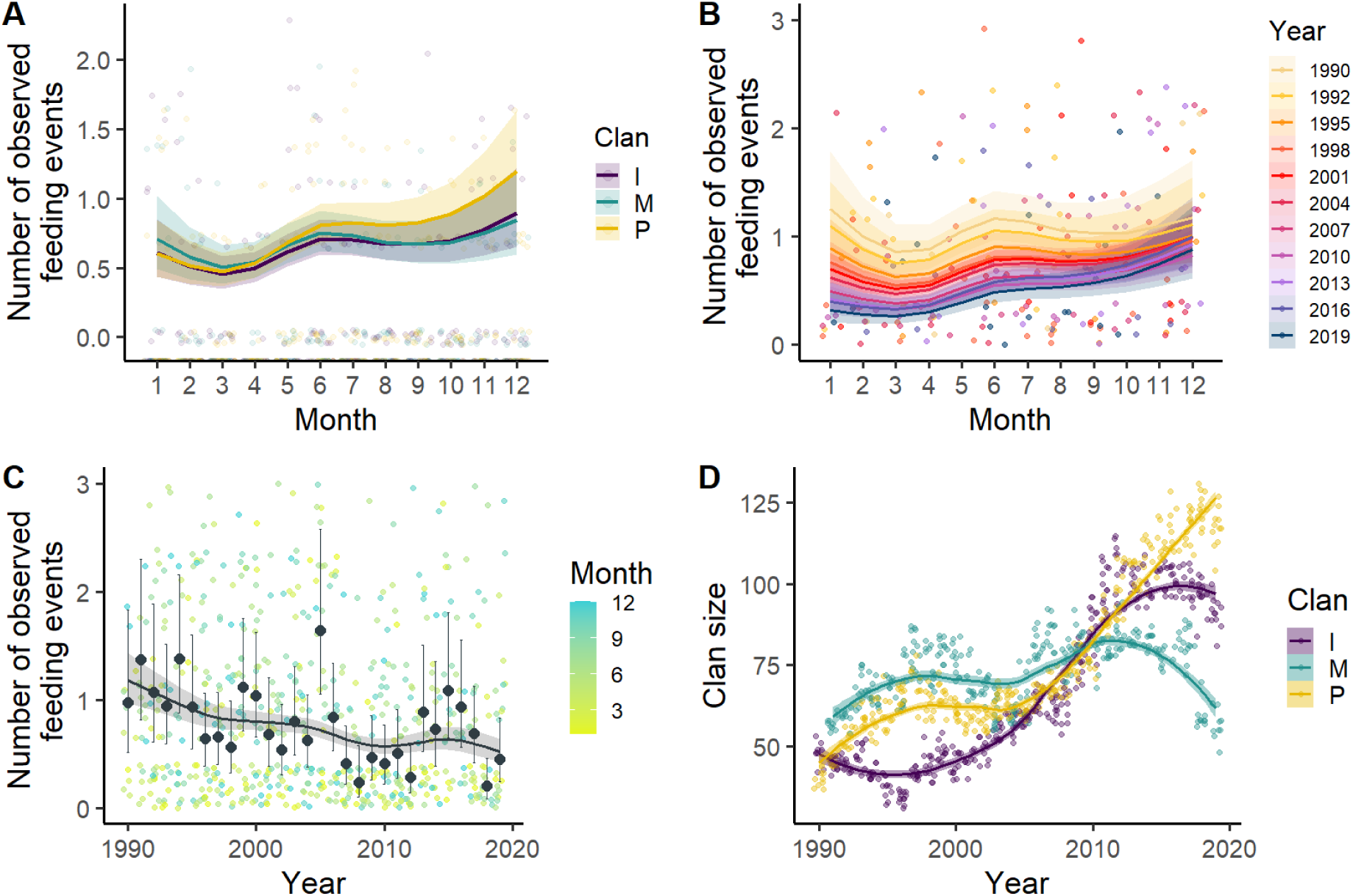
**A)** Annual patterns of the number of observed feeding events for the three clans (estimates produced by the *sMonth*Clan* model. **B)** Changes in annual patterns of the number of observed feeding events over the years (estimates produced by the *sMonth*sYear* model. **C)** Changes in the number of observed feeding events over the years (estimates produced by the *sMonth*sYear* model). **D)** Size of the three different clans over years, where lines and ribbons show the smoothed median of the number of individuals alive in the clan over the years, and dots represent raw data for each clan and each month. For all plots, except D, estimates (solid lines); their 95% confidence intervals (ribbons); and original data (coloured dots) are plotted. Extreme values are not seen on the plots, as only 95% of data is shown.

We then explored how this annual pattern varied over the course of the study period by adding an effect of *Year* to the nonlinear effect of months (*sMonth*; see next paragraph). We chose to use *sMonth*, as it allowed us to substantially reduce the number of parameters while capturing a large amount of variance in annual trends (12.97% for the *sMonth* model versus 4.45% for the *cMonth* model; see Appendix S6).

#### The annual pattern of observed feeding events changed over the years

Changes over the years in annual patterns of the number of observed feeding events matched expectations of the matching hypothesis (Figure 2B). At the beginning of the study period, the annual pattern was characterized by two peaks—one in May-July and another in November-January. This pattern then gradually transitioned into a pattern where the number of observed feeding events was lower in the first half of the year than in the second (estimates from the *sMonth *sYear* model, part of models best predicting trends in *n_feeding_* over the years: ΔAIC = 0 for *sMonth+cYear*, 2.20 for *sMonth*cYear*, 3.36 for *sMonth *sYear,* and 4.71 for *sMonth+sYear;* and ΔAIC = 21.01 for *sMonth+cYear* when compared to the *sMonth* model; see Appendix S6). This shift in annual patterns over the years was primarily driven by a stronger decline in the number of observed feeding events between January and July than in later months and reinforced by a 21% increase in the number of feeding events observed in November and December between 2009 and 2015. Ultimately the number of observed feeding events decreased by 72% in January, 56% in June, and 21% in December from 1990 to 2019 (estimates from the *sMonth*sYear* model; see Appendix S8 for exact values). This flattening of the mid-year peak, along with the stability or slight increase in the number of observed feeding events during late-year months, closely mirrored the previously documented shifts in migratory prey presence within hyaena clan territories (Figure 1, Panel 2; Gicquel et al. 2022).

#### The number of observed feeding events strongly decreased over the years

Overall, the number of observed feeding events decreased by 61% from 1990 to 2019 (estimate from the *sMonth*sYear* model, model selection presented above), despite two periods of stabilization or slight increase (1998-2004 and 2010-2015; Figure 2C). This pattern matched the decline in migratory prey presence previously reported in Gicquel et al. (2022), aligning with the matching hypothesis (Figure 1, Panel 3). All trends over the years were consistent across clans (*sMonth* + *cYear* + *Clan* was the only model including an effect of *Clan* present in models selected to explore annual patterns and trends over the years in *n_feeding_* over the years, ΔAIC = 2.95 among models exploring patterns over the years).

### Change in clan size over the years

Contrary to temporal trends in the number of observed feeding events, the sizes of the Isiaka and Pool clans have increased substantially over years, while the size of the Mamba clan increased until 2010 and then declined steadily until the end of the study period (Figure 2D).

### Relationship between the numbers of observed feeding events and migratory prey presence within hyaena clan territories

As predicted by both the matching and behavioural adjustment hypotheses, the number of feeding events observed was higher in all clans during periods of high migratory prey presence, compared to periods of low or absence of migratory prey presence (estimates from the *Prey_max_* and *Prey_max_* * *Clan* models, ΔAIC = 0 and 2.62 respectively among those exploring an effect of *Prey,* and performing better than models exploring temporal patterns: e.g., ΔAIC= 67.85 for *sMonth+cYear* compared to *Prey_max_* model, see Appendix S9).

## Discussion

We used 30 years of incidental observations of feeding events to investigate how a decline in migratory prey presence within hyaena clan territories influenced the number of hyaena feeding events, and explored trends in hyaena clan size. Temporal trends in the number of observed feeding events closely matched changes in migratory prey presence within hyaena territories (Gicquel et al. 2022b), supporting the matching hypothesis. The number of observed feeding events decreased over the years, with a sharper decline during the first half of the year, than during late-year months. Migratory prey presence was a strong predictor of the number of observed feeding events, with more events observed when migratory prey were abundant. Despite the decline in the number of observed feeding events, two out of the three studied clans increased in size over the study period. Our results, although restricted to feeding events within hyaena clan territories while hyaenas are commuting for much of the year, provide insights into how central-place foragers may respond locally to changes in prey availability, and the extent to which such responses may influence clan size. Few studies have linked local prey availability and feeding dynamics in central place foragers, although territories are a key area where central place foragers should feed when possible.

Among our tested models, those predicting the number of feeding events based on migratory prey presence best explained the data, with a strong and positive relationship that is consistent with established knowledge on hyaena foraging behaviour in the Serengeti ecosystem (Hofer and East 1993a, 1993b, 1993c, Hofer et al. 1995). Annual patterns in the number of observed feeding events closely mirrored patterns of migratory prey presence described by Gicquel et al. (2022) with two peaks: in May-June and November-December (Figure 1, Panel 2). The relatively low number of observed feeding events within clan territories outside these two periods of high migratory prey presence is explained by hyaenas undertaking commuting trips and feeding outside clan territories when migratory prey is locally scarce. Interestingly, estimates of the number of daily feeding events during periods of high migratory prey presence found here were approximately one order of magnitude lower than during early years of the project (∼0.038 [95% CI: 0.033, 0.044] feeding events/day for Prey = 3 in our study vs ∼ 0.18 [95% CI: 0.12, 0.25] feeding events/day for Prey = 3 in Hofer and East (1993a) vs), which likely reflects the general downward trend in the number of feeding events over the 30-year study period we report here.

The annual pattern of observed feeding events changed over the years, and mirrored changes in annual patterns of migratory prey presence in hyaena clan territories, as expected under the matching hypothesis. Notably, the mid-year peak in both the number of observed feeding events and migratory prey presence progressively flattened over time (Figure 1, Panel 2; Figure 2B). This decrease in the mid-year peak likely reflects to some extent changes in climatic factors affecting migratory prey species movement patterns. In particular, increased rainfall during the wet season over the years is suspected to have altered vegetation and soil conditions, and may have enabled the southern plains to sustain herbivores for longer periods, thereby delaying the northward migration of prey (Mtewele et al. 2023). However, despite the decrease in the number of observed feeding events, the sizes of two clans have greatly increased. This increase may be due to a reduction of snaring incidents in more recent years (Hofer et al. 1993, Benhaiem et al. 2023), as well as a potential increase in migratory prey population abundance in the park in the last decades, offering an increased availability of resources in the ecosystem overall (Sinclair et al. 2015). Future analyses based on demographic data are needed to identify the factors and mechanisms responsible for the changes in clan sizes. We expected this absence of a strong positive association between clan size and the number of observed feeding events within territories, as within-territories feeding occurs for a minor part of the year (Hofer and East 1993a), which likely strongly limits its influence on clan size. Nonetheless, the increased clan size may also suggests that hyaenas in these clans have adjusted to the decreasing migratory prey presence in their clan territories over the years. Gicquel et al. (2022) reported an absence of decrease in maternal den attendance over the 30 years of the study period, indicating that hyaenas did not compensate by increasing the duration or frequency of their commuting trips outside their territory. It would be interesting to investigate if non-lactating adult females and immigrant males, which do not have to return regularly to the territories to nurse their offspring, show a similar pattern with a relatively high den attendance over the years. Hyaenas may have found ways within their own territories to cope with reduced migratory prey presence—aside from increasing foraging activity during periods of high migratory prey presence, as temporal patterns in the number of observed feeding events did not support the behavioural adjustment hypothesis.

Among terrestrial carnivorous mammals constrained by territoriality and the use of dens where offspring are sheltered, a common strategy to cope with fluctuating migratory prey availability is to switch the diet to alternative, resident prey when migratory species are absent (Nelson et al. 2012, Elbroch et al. 2013). In some ecosystems, such as Hwange National Park in Zimbabwe, hyaenas display high dietary flexibility, favouring the most abundant and profitable prey (Périquet et al. 2014). In the Maasai Mara National Reserve, which neighbours the Serengeti National Park and is part of the Serengeti ecosystem, hyaenas have been reported to shift their diet toward resident prey to some extent when migratory prey are not available within their territories (Cooper et al. 1999, Holekamp and Dloniak 2011). Here, we observed that, at an annual scale, the proportion of migratory prey among the feeding events observed was slightly lower in February and March (see Appendix S5), when migratory herbivores are in the southern area of their migration to give birth (Wilmshurst et al. 1999, Mtewele et al. 2023). Yet, despite this seasonal trend, hyaenas did not appear to have increased the proportion of resident prey in their diet over the years (see Appendix S5). The apparent absence of a shift to resident prey we observe here is consistent with the constraints imposed by low resident prey densities within hyaena clan territories, which are insufficient to sustain the large hyaena clan sizes observed in the Serengeti (Hofer and East 1993a, Hofer et al. 1995). The consistently high proportion of migratory prey among observed feeding events aligns with the findings in Hofer and East (1993a) who reported that about 70% of hyaena diet consisted of migratory prey, namely blue wildebeest (54%) and Thomson’s gazelles (15%).

As clan size increased despite the decreasing number of observed feeding events, hyaenas may have relied on alternative resources beyond migratory and resident prey. In other ecosystems, hyaenas are known to use anthropogenic food subsidies, such as food waste produced in human settlements and refuse pits, which can influence hyaena space use and allow them to reach higher densities than populations not benefiting from these resources (Kolowski and Holekamp 2008, Yirga et al. 2012). In recent years, adults and subadults of one of the study clans have occasionally been observed entering a tourist camp, and are suspected to enter the camp to look for trash and human food remains (S. Metzger, personal communication). The waste produced in touristic areas offer a resource that is predictable in space and time, for which hyaenas only need to perform a round trip and to return to their den, without the need to extend their commuting trips. However, given the low number of observations of such behaviour and the large amount of resources required to sustain the three large hyaena clans studied, we do not expect human food waste to constitute a main food source. Nevertheless, future studies could apply for an authorization to deploy camera traps at dump sites in the Seronera area to investigate if hyaenas do come feed at dump sites and whether this a regularly-used strategy.

It is worth noting that we could not fully account for the detection probability of feeding events, which may have influenced our results. Although we included the number of monitoring sessions conducted in a given month (which are the direct conditions for incidental feeding events observations) as an offset in the model, other sources of variation in observation effort were not considered. These include observer-related factors, such as the number, identity, or experience of observers, which could have affected detection rates. Environmental factors such as rainfall, which influence grass height and visibility, were also not accounted for. Tall grass can reduce the detectability of feeding events, especially when few individuals are present, and heavy rains limit off-road driving, making feeding events far from roads more difficult to detect. In addition, a potential bias could arise from prey size. Smaller prey, such as Thomson’s gazelles, or hares (*Lepus capensis*), are typically consumed much more rapidly than large prey, and small carcasses attract less hyaenas individuals to feed (Moleón et al. 2015), making them less likely to be detected. However, we do not expect that missing this type of small prey would considerably modify the patterns detected, as they are not expected to constitute a substantial part of hyaena diet due to their densities being too low to sustain large hyaena clan sizes (Hofer and East 1993a, Hofer et al. 1995). Among small prey, Thomson’s gazelles are an exception, as they are relatively abundant and may make up a substantial part of hyaena diet. However, the proportion of feeding events involving this species did not show a consistent decline over time, suggesting that reduced detection of Thompson’s gazelle feeding event is unlikely to explain the observed decrease in total feeding events over the years (Appendix S10). Moreover, observation effort was consistently high, with many monitoring sessions conducted each month over the 30-year period.

Until now, we have interpreted the decline in migratory prey within hyena territories as a true decrease in local prey availability. However, it is also possible that changes in prey spatial distribution could also lead to the results we observe without true changes in hyaena foraging strategy. Increased human activity can influence ungulate spatial distribution (Romero et al. 2024). To access the clans, observers consistently started from the more tourist-frequented area around Seronera, where the Seronera village, the Serengeti Wildlife Research Center and campsites and lodges are located. Yet wildebeest, the main prey of hyaena in the Serengeti ecosystem, may have reduced their occupancy of areas frequented by humans over the years (*Serengeti Biodiversity Program Annual Report 2023-2024* 2025), which coincided with an increase in the number of visitors in the park over the years (TANAPA data provided in Mbisse 2021). We can therefore describe a potential scenario in which the spatial distribution of migratory prey around the three hyaena clan territories have shifted over the years, such that they would less often detected by observers even though large herds actually still occurred in the area. Since the majority of feeding events occur where migratory prey are present, this could also explain the decline in the number of observed feeding events over the years. From a hyaena point of view, no real changes in resources distribution (in the clan territories) have emerged in the last decades under this hypothesis, thereby explaining that during the same period there was no decrease in 1) maternal den attendance, i.e. no increase in long-distance commuting trips (Gicquel et al. 2022b), 2) cub survival and recruitment (White et al. 2025) and 3) clan size (this study). To assess whether migratory prey remain present in hyaena territories but in areas not monitored by observers, transect surveys similar to those conducted by Hofer and East (1993a) could be employed to estimate prey abundance outside the primary study area and investigate potential shifts in prey distribution in the Seronera area. However, the methodology should be adapted to the landscape characteristics of the different areas (e.g. bush thickness).

We studied the within-territory correlation between migratory prey presence and feeding events in the territories of a central-place forager. These interactions only capture a partial picture of how prey availability influences the foraging behaviour and clan size of central-place foragers. However, even though within-territory feeding occurs only during a minor part of the year in our study area, hyaenas still have to feed during such periods. Given that hyaenas do not appear to have increased their commuting effort (Gicquel et al. 2022b), it is plausible that they modified their behaviour locally to adjust to reduced migratory prey presence. Understanding these potential local-scale response would offer broader insights into how central-place foragers adapt to spatial and temporal shifts in prey availability. Beyond the Serengeti’s hyaenas, many predators that rely on migratory prey are increasingly struggling to track the spatiotemporal distribution of their prey, as climate change alters migration patterns (Durant et al. 2007, Carroll et al. 2024). Understanding the consequences of these spatial and phenological changes, and the strategies predators use to adjust, is crucial for predicting predators future in a warming world where climate change is projected to persist and intensify (Intergovernmental Panel on Climate Change (IPCC) 2023).

## Acknowledgments

We thank the Commission for Science and Technology of Tanzania (COSTECH) for permission to conduct research, and the Tanzania Wildlife Research Institute (TAWIRI) and Tanzania National Parks (TANAPA) for their support. We thank the Deutsche Forschungsgemeinschaft (grants EA 5/3-1, KR 4266/2-1, DFG-Grako 1121, DFG-Grako 2046), Leibniz-Gemeinschaft (grant SAW-2018-IZW-3-EpiRank), Leibniz-Institute for Zoo and Wildlife Research, Fritz-Thyssen-Stiftung, and Stifterverband der deutschen.

## Author contribution

MS, MG, SB and SC designed the study. MS extracted data and performed the analyses, with assistance from SC, SB and MG. MS drafted the manuscript with contribution by SB. MLE, SM, HH and SB collected the field data. MLE and HH coordinated the long-term project with assistance from JN. All authors provided input on the manuscript and edited the final draft.

## Conflict of Interest

The authors declare no conflict of interest.

## Notes

### Competing Interest Statement

The authors have declared no competing interest.

## References

Anholt, B. R., and E. E. Werner. 1998. Predictable changes in predation mortality as a consequence of changes in food availability and predation risk. Evolutionary Ecology 12:729–738.

Benhaiem, S., S. Kaidatzi, H. Hofer, and M. L. East. 2023. Long-term reproductive costs of snare injuries in a keystone terrestrial by-catch species. Animal Conservation 26:61–71.

Boone, R. B., S. J. Thirgood, and J. G. C. Hopcraft. 2006. Serengeti Wildebeest Migratory Patterns Modeled from Rainfall and New Vegetation Growth. Ecology 87:1987–1994.

Burnham, K. P., and D. R. Anderson. 2010. Model selection and multimodel inference: a practical information-theoretic approach. 2. ed. Springer, New York, NY.

Carroll, G., B. Abrahms, S. Brodie, and M. A. Cimino. 2024. Spatial match–mismatch between predators and prey under climate change. Nature Ecology & Evolution:1–9.

Cooper, S. M., K. E. Holekamp, and L. Smale. 1999. A seasonal feast: long-term analysis of feeding behaviour in the spotted hyaena (Crocuta crocuta). African Journal of Ecology 37:149–160.

Durant, J. M., D. Ø. Hjermann, G. Ottersen, and N. C. Stenseth. 2007. Climate and the match or mismatch between predator requirements and resource availability. Climate Research 33:271–283.

Elbroch, L. M., P. E. Lendrum, J. Newby, H. Quigley, and D. Craighead. 2013. Seasonal Foraging Ecology of Non-Migratory Cougars in a System with Migrating Prey. PLOS ONE 8:e83375.

Frank, L. G. 1986a. Social organization of the spotted hyaena *Crocuta crocuta*. II. Dominance and reproduction. Animal Behaviour 34:1510–1527.

Frank, L. G. 1986b. Social organization of the spotted hyaena (*Crocuta crocuta*). I. Demography. Animal Behaviour 34:1500–1509.

Frank, L. G., S. E. Glickman, and P. Licht. 1991. Fatal Sibling Aggression, Precocial Development, and Androgens in Neonatal Spotted Hyenas. Science 252:702–704.

Gicquel, M., M. L. East, H. Hofer, and S. Benhaiem. 2022a. Early-life adversity predicts performance and fitness in a wild social carnivore. Journal of Animal Ecology 91:2074–2086.

Gicquel, M., M. L. East, H. Hofer, S. Cubaynes, and S. Benhaiem. 2022b. Climate change does not decouple interactions between a central-place-foraging predator and its migratory prey. Ecosphere 13:e4012.

Hartig, F. 2024.DHARMa: Residual Diagnostics for Hierarchical (Multi-Level / Mixed) Regression Models.

Hastie, T. 2024. gam: Generalized Additive Models.

Hetem, R. S., A. Fuller, S. K. Maloney, and D. Mitchell. 2014. Responses of large mammals to climate change. Temperature 1:115–127.

Hofer, H., M. East, H. Hofer, and M. East. 1995. Population dynamics, population size, and the commuting system of Serengeti spotted hyenas. Sinclair, A R E, Arcese, P Serengeti II:Dynamics, management, and conservation of an ecosystem 332–363.

Hofer, H., and M. L. East. 1993a. The commuting system of Serengeti spotted hyaenas: how a predator copes with migratory prey. I. Social organization. Animal Behaviour 46:547–557.

Hofer, H., and M. L. East. 1993b. The commuting system of Serengeti spotted hyaenas: how a predator copes with migratory prey. II. Intrusion pressure and commuters’ space use. Animal Behaviour 46:559–574.

Hofer, H., and M. L. East. 1993c. The commuting system of Serengeti spotted hyaenas: how a predator copes with migratory prey. III. Attendance and maternal care. Animal Behaviour 46:575–589.

Hofer, H., M. L. East, and K. L. Campbell. 1993. Snares, commuting hyaenas and migratory herbivores: humans as predators in the Serengeti. Symposia of the Zoological Society of London 65.

Holekamp, K., and S. Dloniak. 2011. Intraspecific Variation in the Behavioral Ecology of a Tropical Carnivore, the Spotted Hyena. Advances in the Study of Behavior 42:189.

Höner, O. P., B. Wachter, M. L. East, and H. Hofer. 2002. The response of spotted hyaenas to long-term changes in prey populations: functional response and interspecific kleptoparasitism. Journal of Animal Ecology 71:236–246.

Höner, O. P., B. Wachter, M. L. East, V. A. Runyoro, and H. Hofer. 2005. The effect of prey abundance and foraging tactics on the population dynamics of a social, territorial carnivore, the spotted hyena. Oikos 108:544–554.

Iler, A. M., P. J. CaraDonna, J. R. K. Forrest, and E. Post. 2021. Demographic Consequences of Phenological Shifts in Response to Climate Change. Annual Review of Ecology, Evolution, and Systematics 52:221–245.

Intergovernmental Panel on Climate Change (IPCC). 2023. Climate Change 2021 – The Physical Science Basis: Working Group I Contribution to the Sixth Assessment Report of the Intergovernmental Panel on Climate Change. Cambridge University Press, Cambridge.

Jeanniard-du-Dot, T., A. W. Trites, J. P. Y. Arnould, and C. Guinet. 2017. Reproductive success is energetically linked to foraging efficiency in Antarctic fur seals. PLOS ONE 12:e0174001.

Kolowski, J. M., and K. E. Holekamp. 2008. Effects of an open refuse pit on space use patterns of spotted hyenas. African Journal of Ecology 46:341–349.

Labadie, G., C. Hardy, Y. Boulanger, V. Vanlandeghem, M. Hebblewhite, and D. Fortin. 2023. Global change risks a threatened species due to alteration of predator–prey dynamics. Ecosphere 14:e4485.

Lecomte, N., G. Gauthier, and J.-F. Giroux. 2009. A link between water availability and nesting success mediated by predator-prey interactions in the Arctic. Ecology 90:465–475.

Lüdecke, D. 2018. ggeffects: Tidy Data Frames of Marginal Effects from Regression Models._, *3*(26), 772. doi:10.21105/joss.00772 <10.21105/joss.00772>. Journal of Open Source Software 26:772.

Mahony, J. 2020. Modelling the impact of climate change on the wildebeest of the Serengeti-Mara ecosystem. http://purl.org/dc/dcmitype/Text, University of Oxford.

Mbisse, R. Z. 2021. Forecasting the tourist arrivals at Serengeti national park in Tanzania. University of Dodoma, Tanzania.

Mills, L., and F. Knowlton. 1991. Coyote Space Use in Relation to Prey Abundance. Canadian Journal of Zoology-Revue Canadienne De Zoologie 69:1516–1521.

Moleón, M., J. A. Sánchez-Zapata, E. Sebastián-González, and N. Owen-Smith. 2015. Carcass size shapes the structure and functioning of an African scavenging assemblage. Oikos 124:1391–1403.

Morin, A., S. Chamaillé-Jammes, and M. Valeix. 2021. Climate Effects on Prey Vulnerability Modify Expectations of Predator Responses to Short- and Long-Term Climate Fluctuations. Frontiers in Ecology and Evolution 8:601202.

Mtewele, Z. F., G. Jia, and X. Xu. 2023. Serengeti–Masai Mara ecosystem dynamics inferred from rainfall extremes. Environmental Research Letters 18:114026.

Naciri, M., A. Planillo, M. Gicquel, M. L. East, H. Hofer, S. Metzger, and S. Benhaiem. 2023. Three decades of wildlife-vehicle collisions in a protected area: Main roads and long-distance commuting trips to migratory prey increase spotted hyena roadkills in the Serengeti. Biological Conservation 279:109950.

Nelson, A. A., M. J. Kauffman, A. D. Middleton, M. D. Jimenez, D. E. McWhirter, J. Barber, and K. Gerow. 2012. Elk migration patterns and human activity influence wolf habitat use in the Greater Yellowstone Ecosystem. Ecological Applications 22:2293–2307.

Ogutu, J. O., G. S. Bartzke, S. Mukhopadhyay, H. T. Dublin, J. S. Senteu, D. Gikungu, I. Obara, and H.-P. Piepho. 2024. Trends and cycles in rainfall, temperature, NDVI, IOD and SOI in the Mara-Serengeti: Insights for biodiversity conservation. PLOS Climate 3:e0000388.

Périquet, S., H. Fritz, and E. Revilla. 2014. The Lion King and the Hyaena Queen: large carnivore interactions and coexistence. Biological Reviews 90:1197–1214.

R Core Team. 2022. R: A Lnguage and Environment for Statistical Computing. R Foundation for Statistical Computing, Vienna, Austria.

Romero, A., B. J. O’Neill, K. Rauch, and A. Roscoe. 2024. How African Ungulates Respond to Tourist Vehicles in Kruger National Park. African Journal of Ecology 62:e13335.

Ronconi, R. A., and A. E. Burger. 2008. Limited foraging flexibility: increased foraging effort by a marine predator does not buffer against scarce prey. Marine Ecology Progress Series 366:245–258.

Roth, J. 2003. Variability in marine resources affects arctic fox population dynamics. Journal of Animal Ecology 72:668–676.

Schmidt, K. 2008. Behavioural and spatial adaptation of the Eurasian lynx to a decline in prey availability. Acta Theriologica 53:1–16.

Schmidt, N. M., R. A. Ims, T. T. Høye, O. Gilg, L. H. Hansen, J. Hansen, M. Lund, E. Fuglei, M. C. Forchhammer, and B. Sittler. 2012. Response of an arctic predator guild to collapsing lemming cycles. Proceedings of the Royal Society B: Biological Sciences 279:4417–4422.

Serengeti Biodiversity Program Annual Report 2023-2024. 2025. . Serengeti Biodiversity Program, 4th Greater Serengeti-Mara Ecosystem forum.

Sinclair, A. R. E., K. L. Metzger, S. A. R. Mduma, and J. M. Fryxell. 2015. Serengeti IV: Sustaining Biodiversity in a Coupled Human-Natural System. University of Chicago Press.

Stewart, J. D., J. W. Durban, H. Fearnbach, L. G. Barrett-Lennard, P. K. Casler, E. J. Ward, and D. R. Dapp. 2021. Survival of the fattest: linking body condition to prey availability and survivorship of killer whales. Ecosphere 12:e03660.

Strauss, E. D., F. H. Jensen, A. S. Gersick, M. Thomas, K. E. Holekamp, and A. Strandburg-Peshkin. 2024. Daily ranging and den usage patterns structure the spatiotemporal properties of social encounters in spotted hyenas. Behavioral Ecology and Sociobiology 78:45.

Tablado, Z., P. Fauchald, G. Mabille, A. Stien, and T. Tveraa. 2014. Environmental variation as a driver of predator-prey interactions. ECOSPHERE 5:164.

Venables, W. N., B. D. Ripley, and W. N. Venables. 2002. Modern applied statistics with S. 4th ed. Springer, New York.

Walton, L. R., H. D. Cluff, P. C. Paquet, and M. A. Ramsay. 2001. Movement Patterns of Barren-Ground Wolves in the Central Canadian Arctic. Journal of Mammalogy 82:867–876.

White, E., J. Louvrier, L. Bailey, E. Davidian, M. East, H. Hofer, B. Wachter, S. Benhaiem, O. Höner, and V. Radchuk. 2025. Resilience of a long-lived mammal: time and demographic structure matter. Authorea.

Wickham, H. 2016. ggplot2: elegant graphics for data analysis. Second edition. Springer, Switzerland.

Wilmshurst, J. F., J. M. Fryxell, B. P. Farm, A. Sinclair, and C. P. Henschel. 1999. Spatial distribution of Serengeti wildebeest in relation to resources. Canadian Journal of Zoology 77:1223–1232.

Yirga, G., H. H. De Iongh, H. Leirs, K. Gebrihiwot, J. Deckers, and H. Bauer. 2012. Adaptability of large carnivores to changing anthropogenic food sources: diet change of spotted hyena (*Crocuta crocuta*) during Christian fasting period in northern Ethiopia. The Journal of Animal Ecology 81:1052–1055.

