## supplementary material for "How did spotted hyaenas respond to decreased prey availability in their clan territories over the last decades?"

**Appendix S1: Proportion of each index of prey abundance within the three spotted hyaena clan territories over the years**

**Table S1:** Yearly proportions of each index of prey abundance (as defined in Hofer & East 1993a) within the three spotted hyaena clan territories.

| | **Migratory prey** | | |
|---|---|---|---|
| **Year** | **Abundant (Prey = 3)** | **Low (Prey = 2)** | **Absent (Prey = 1)** |
| 1990 | 0.20 | 0.24 | 0.56 |
| 1991 | 0.07 | 0.60 | 0.33 |
| 1992 | 0.18 | 0.22 | 0.60 |
| 1993 | 0.03 | 0.61 | 0.36 |
| 1994 | 0.10 | 0.52 | 0.38 |
| 1995 | 0.15 | 0.44 | 0.40 |
| 1996 | 0.08 | 0.42 | 0.49 |
| 1997 | 0.15 | 0.50 | 0.35 |
| 1998 | 0.15 | 0.46 | 0.39 |
| 1999 | 0.20 | 0.57 | 0.23 |
| 2000 | 0.10 | 0.71 | 0.20 |
| 2001 | 0.04 | 0.49 | 0.47 |
| 2002 | 0.04 | 0.65 | 0.31 |
| 2003 | 0.14 | 0.63 | 0.23 |
| 2004 | 0.04 | 0.55 | 0.41 |
| 2005 | 0.09 | 0.53 | 0.38 |
| 2006 | 0.04 | 0.47 | 0.49 |
| 2007 | 0.11 | 0.58 | 0.31 |
| 2008 | 0.07 | 0.44 | 0.49 |
| 2009 | 0.17 | 0.49 | 0.34 |
| 2010 | 0.03 | 0.64 | 0.33 |
| 2011 | 0.05 | 0.47 | 0.48 |
| 2012 | 0.03 | 0.47 | 0.50 |
| 2013 | 0.02 | 0.45 | 0.53 |
| 2014 | 0.09 | 0.33 | 0.57 |
| 2015 | 0.04 | 0.59 | 0.37 |
| 2016 | 0.07 | 0.61 | 0.32 |
| 2017 | 0.12 | 0.57 | 0.30 |
| 2018 | 0.05 | 0.38 | 0.57 |
| 2019 | 0.09 | 0.45 | 0.46 |

**Appendix S2: List of variables and their abbreviations**

**Table S1:** Overview of the variables used in the statistical analysis and their abbreviations

| Abbreviation | Definition | Type of variable |
|---|---|---|
| $n_{feeding}$ | Number of monthly incidental observations of feeding events in each hyaena clan territory | Response |
| *Prey_type* | Type of prey consumed (2 levels: 0 = resident; 1 = migratory) | |
| *Prey* | Migratory prey presence within hyaena clan territories (3 levels: 1 = absence of migratory prey ; 2 = low abundance of migratory prey; 3 = high abundance of migratory prey, ) | Predictor |
| $Prey_{mode}$ | Most common value of *Prey* recorded a given month for a given clan | |
| $Prey_{max}$ | Maximum recorded value of *Prey* recorded a given month for a given clan | |
| *Clan* | Clan affiliation (3 levels: I, M, P) | |
| *Month* | Month during which observations were made (12 levels; from 1 = January, to 12 = December) | |
| *qMonth* | Quadratic effect of the variable *Month* | |
| *cMonth* | Categorical effect of *Month* | |
| *sMonth* | Nonlinear effect of *Month* | |
| *Year* | Year during which observations were made (30 levels, from 1990 to 2019) | |
| *lYear* | Linear effect of *Year* | |
| *qYear* | Quadratic effect of *Year* | |
| *cYear* | Categorical effect of *Year* | |
| *sYear* | Nonlinear effect of *Year* | |
| *rYear* | Random effect of *Year* | |
| $n_{monitored}$ | Monthly number of days monitored in each clan | Offset |

*Note:* Variables names are given in italics.

### Appendix S3: Additional information about the cause of death of prey

In total, 777 feeding events have been observed (284 for Isiaka, 203 for Mamba and 290 for Pool). Out of these, we had access to the type of prey being consumed for 732 feeding events, (260 for Isiaka, 198 for Mamba and 274 for Pool). The prey consumed was a migratory prey for 601 feeding events recorded (216 for Isiaka, 162 for Mamba and 223 for Pool). Among the carcasses observed belonging to a migratory prey, 325 feeding events were that of the blue wildebeest *Connochaetes taurinus* (130 for Isiaka, 83 for Mamba and 112 for Pool), 167 were that of a Thompson's gazelle *Eudorcas thomsonii* (54 for Isiaka, 48 for Mamba and 65 for Pool) and 109 that of a plain's zebra *Equus quagga* (32 for Isiaka, 31 for Mamba and 46 for Pool). The two non-migratory prey species that were the most consumed were the buffalo (*Syncerus caffer* ; 54 feeding events in total, 13 for Isiaka and Mamba each and 28 for Pool) and the warthog (*Aper aethiopicus* ; 24 feeding events in total, 4 for Isiaka and 10 for Mamba and Pool each). The other species of resident prey observed were Grant gazelle (*Nanger granti* ; n=10), topi (*Damaliscus lunatus jimela* ; n=8), kongoni (*Alcelaphus buselaphus* ; n=1), giraffe (*Giraffa camelopardalis* ; n=13), impala (*Aepyceros melampus* ; n=2), reedbuck (*Redunca spp* ; n=4), elephant (*Loxodonta africana* ; n=3) and hippopotamus (*Hippopotamus amphibus* ; n=3), olive baboon (*Papio anubis*, n= 1), Cheetah (*Acinonyx jubatus*, n=1), dik-dik (*Madoqua kirkii* ; n=2), hare (*Lepus capensis*, n=1), lion (*Panthera leo*, n=1), python (*Python sebae*, n=1), wildcat (*Felis lybica*, n=1).

### Appendix S4 : Goodness of fit of the global model explaining the number of observed feeding events ($n_{feeding}$)

Over the 30 years of the study period, 360 months elapsed. Out of theses 360 months, 135 months in Isiaka, 119 in Mamba and 142 in Pool have non-zeros values for $n_{feeding}$, meaning that at least one feeding event was observed, and on average 0.79 ($\sigma = 1.62$) feeding events per clan have been observed each month since January 1990. The maximum number of feeding events recorded during a month was 10 in the clan Isiaka and 7 for Mamba and Pool. Because of the excess of zeros and the high variance compared to the mean, the response variable deviated strongly from a Poisson distribution. When using a Poisson distribution in Generalized Additive Models that estimated $n_{feeding}$, the overdispersion test from the DHARMa package (Hartig 2024) indicated a poor fit: the statistic of the test was significant ($p$=0) and the overdispersion parameter (1.55) was close to 1.5 (Figure S1A). This issue was solved when using a negative binomial distribution instead (Figure S1B): the test was no longer significant ($p$=0.552) and the dispersion parameter fell below 1.5 (equal to 0.91).

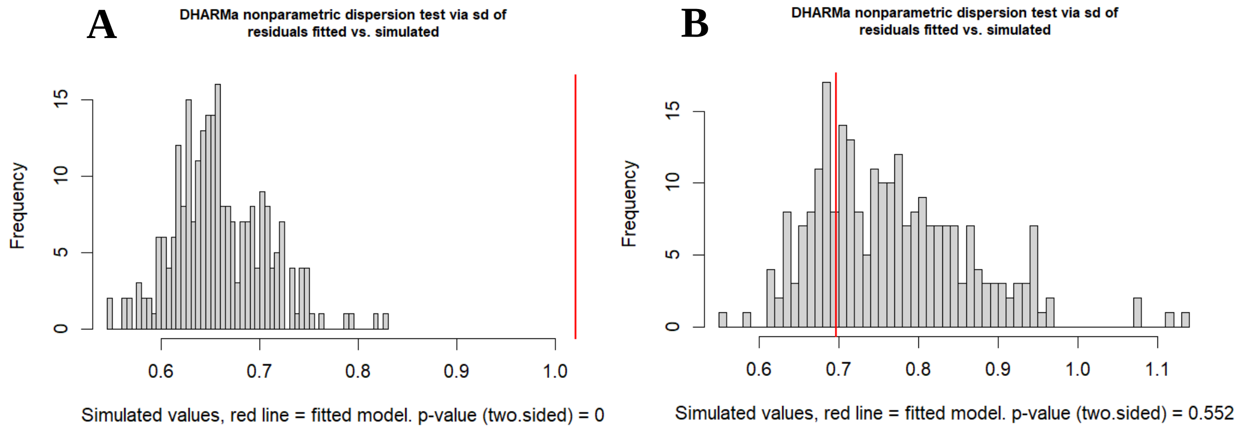


**Figure S1:** Overdispersion test using the DHARMa package (Hartig 2024) and a model *cMonth*Year*Clan* (see Appendix S2, Table S1 for list of variable abbreviations and their meaning). **A)** Using a Poisson distribution. **B)** Using a negative binomial distribution.

Hartig, F. 2024. DHARMa: Residual Diagnostics for Hierarchical (Multi-Level / Mixed) Regression Models.

**Appendix S5: Temporal variation in the type of prey consumed (*Prey_type*)**

Since resident prey occur at low densities and are insufficient to sustain hyaena clans in the Serengeti in the absence of migratory prey, we did not expect a shift of diet to resident prey to be a possible strategy for hyaenas to cope with reduced migratory prey presence in their clan territories. Hence, we did not expect to observe changes in the proportion of migratory prey among the observed feeding events over years but aimed to verify this formally.

*Statistical analysis*

To verify that hyaenas could not shift their diet to alternative prey following the reduction in migratory prey presence (*Prey;* see Appendix S2, Table S1 for list of variables abbreviations and their meaning) over the years, we investigated how *Prey_type* (value 1 if the prey is migratory and 0 is the prey is resident, as explained in main text) varied over the 30-year study period using Generalized Linear Models and Generalized Additive Models. We conducted the same analysis as the one presented in the main text for $n_{feeding}$ without including an offset, as we did not expect the type of prey consumed to depend on observation effort.

*Results*

At an annual scale, the proportion of migratory prey consumed during observed feeding events was high year-round. It peaked in June and November while being lowest in February-March (*Prey_type* = 0.91 [95%CI: 0.83, 0.95] in June and 0.62 [95%CI: 0.49, 0.73] in February; estimates from the *cMonth* and *sMonth* models, ΔAIC = 0 and 2.31, respectively, among models predicting annual trends in *Prey_type*; see Table S1 below for model selection; Figure S1A). As expected, hyaenas did not increase the relative proportion of resident prey in feeding events over the years, as *Prey_type* remained consistently high (estimates from the model *sMonth*cYear*, ΔAIC = 0; Figure S1B). Temporal trends were similar across clans, although the relative proportion of migratory prey was slightly higher in Mamba — 4% and 7% higher than Pool and Isiaka, respectively (estimates from the *sMonth* * *cYear* + *Clan* model, ΔAIC = 3.35 among models exploring temporal trends). The proportion of migratory prey consumed during feeding events was particularly high during periods of high migratory prey presence in the clan territories ($Prey_{max}$ model, ΔAIC = 3.59 in the overall model selection), and this was consistent across clans (no *Clan* effect in models selected based on AIC).

**Table S1:** Model selection for investigating temporal trends in the proportion of migratory prey among observed feeding events (*Prey_type*).

| Explanatory variable | $R^2$ | AIC | ΔAIC |
|---|---|---|---|
| **sMonth * cYear** | **27.11** | **627.33** | **0** |
| **sMonth * cYear + Clan** | **27.21** | **630.67** | **3.35** |
| **$Prey_{max}$** | **9.14** | **630.92** | **3.59** |
| $Prey_{max}$ + Clan | 9.17 | 634.76 | 7.43 |
| $Prey_{max}$ * Clan | 9.51 | 640.39 | 13.06 |
| $Prey_{mode}$ | 5.78 | 654.03 | 26.70 |
| $Prey_{mode}$ + Clan | 5.85 | 657.54 | 30.21 |
| $Prey_{mode}$ * Clan | 6.93 | 658.11 | 30.79 |
| sMonth + cYear | 14.10 | 658.84 | 31.51 |
| sMonth + cYear + Clan | 14.10 | 662.82 | 35.50 |
| sMonth * Year * Clan | 7.64 | 665.29 | 37.96 |
| sMonth * qYear * Clan | 7.64 | 665.29 | 37.96 |
| sMonth + Year * Clan | 5.86 | 667.52 | 40.20 |
| sMonth * sYear | 5.80 | 667.93 | 40.60 |
| sMonth * Year | 4.90 | 668.12 | 40.80 |
| sMonth * qYear | 4.90 | 668.12 | 40.80 |
| sMonth * sYear * Clan | 7.96 | 669.04 | 41.71 |
| sMonth + qYear * Clan | 6.20 | 669.16 | 41.84 |
| sMonth + Year | 4.41 | 669.44 | 42.11 |
| sMonth + qYear | 4.41 | 669.44 | 42.11 |
| sMonth + sYear | 5.21 | 669.95 | 42.62 |
| sMonth * sYear + Clan | 5.91 | 671.14 | 43.81 |
| sMonth + sYear * Clan | 6.73 | 671.52 | 44.20 |
| sMonth * Year + Clan | 4.95 | 671.76 | 44.44 |
| sMonth * qYear + Clan | 4.95 | 671.76 | 44.44 |
| sMonth + sYear + Clan | 5.34 | 673.05 | 45.72 |
| sMonth + Year + Clan | 4.47 | 673.06 | 45.73 |
| sMonth + qYear + Clan | 4.47 | 673.06 | 45.73 |
| cMonth | 5.41 | 674.58 | 47.25 |
| sMonth | 3.04 | 676.88 | 49.56 |
| cMonth + Clan | 5.56 | 677.57 | 50.24 |
| cMonth * Clan | 11.79 | 678.75 | 51.42 |
| sMonth + Clan | 3.14 | 680.19 | 52.86 |
| sMonth * Clan | 3.29 | 683.20 | 55.87 |
| qMonth | 1.02 | 686.82 | 59.50 |
| qMonth + Clan | 1.09 | 690.30 | 62.98 |
| qMonth * Clan | 1.87 | 692.98 | 65.65 |
| sMonth + cYear * Clan | 25.66 | 699.31 | 71.99 |
| sMonth + cYear * Clan | 44.89 | 719.06 | 91.74 |

*Note:* The abbreviations *l* indicates a linear effect of the variable, *c* a categorical, *s* a non-linear effect and *q* a quadratic effect. The best equivalent models are in bold.

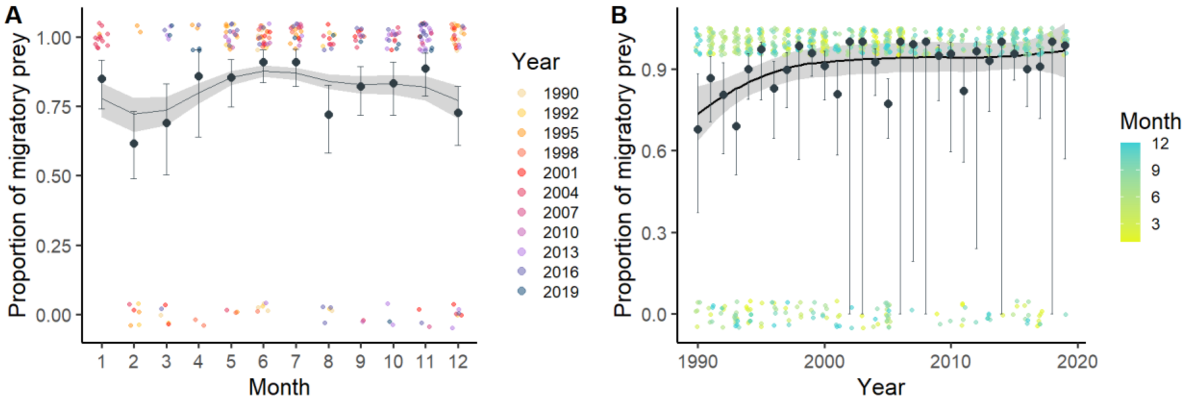


**Figure S1:** Variation of the proportion of migratory prey among observed feeding events (*Prey_type*; on the y-axis, 0 indicates that the prey consumed was a resident species, and 1 that it was a migratory species) over months and years. **A)** Annual pattern of *Prey_type* estimated from a model with categorical effect of month (*cMonth*; points and error bars) and a model with a non-linear effect of month (*sMonth*; line and ribbon). **B)** Patterns over years of *Prey_type* estimated from a model with a categorical effect of *Year* (model *sMonth+cYear)*. Points and error bars represent model estimates and their 95% confidence interval. Line and ribbon represent the smooth of the trend. For both panels, coloured dots represent raw data, , that were jittered along the x and y-axis to improve visibility.

**Appendix S6: Model selection for investigating trends in the number of observed feeding events.**

**Table S1:** Model selection for investigating the temporal trend in $n_{feeding}$ and its links to Prey and Clan.

| Explanatory variable | $R^2$ | AIC | ΔAIC |
|---|---|---|---|
| **Prey$_{max}$** | **12,11** | **2136,32** | **0** |
| **Prey$_{max}$ + Clan** | **12,33** | **2138,14** | **1,82** |
| **Prey$_{max}$ * Clan** | **12,98** | **2139,94** | **3,62** |
| Prey$_{mode}$ | 8,23 | 2172,94 | 36,62 |
| Prey$_{mode}$ + Clan | 8,41 | 2175,26 | 38,94 |
| Prey$_{mode}$ * Clan | 8,65 | 2181,00 | 44,68 |
| sMonth + cYear | 21,26 | 2204,17 | 67,85 |
| sMonth + rYear | 21,26 | 2206,17 | 69,85 |
| sMonth * cYear | 27,12 | 2206,37 | 70,05 |
| sMonth + cYear + Clan | 21,37 | 2207,12 | 70,80 |
| sMonth * sYear | 15,87 | 2207,53 | 71,21 |
| sMonth + sYear | 15,52 | 2208,88 | 72,56 |
| sMonth + rYear + Clan | 21,37 | 2209,12 | 72,80 |
| sMonth * cYear + Clan | 27,21 | 2209,45 | 73,13 |
| sMonth * qYear | 15,39 | 2210,10 | 73,78 |
| sMonth * Year | 14,95 | 2210,29 | 73,97 |
| sMonth * sYear + Clan | 15,99 | 2210,34 | 74,02 |
| sMonth + sYear + Clan | 15,64 | 2211,68 | 75,36 |
| sMonth + rYear * Clan | 21,51 | 2211,75 | 75,43 |
| sMonth + Year | 14,55 | 2212,11 | 75,79 |
| sMonth * qYear * Clan | 17,63 | 2212,73 | 76,41 |
| sMonth + sYear * Clan | 16,37 | 2212,78 | 76,46 |
| sMonth * qYear + Clan | 15,51 | 2212,90 | 76,58 |
| sMonth * Year + Clan | 15,07 | 2213,15 | 76,83 |
| sMonth + qYear | 14,63 | 2213,29 | 76,97 |
| sMonth * sYear * Clan | 16,86 | 2214,06 | 77,75 |
| sMonth + Year + Clan | 14,67 | 2214,95 | 78,63 |
| sMonth + qYear * Clan | 16,10 | 2215,29 | 78,97 |
| sMonth + Year * Clan | 15,45 | 2215,55 | 79,23 |
| sMonth + qYear + Clan | 14,77 | 2216,02 | 79,70 |
| sMonth * Year * Clan | 16,00 | 2216,24 | 79,92 |
| sMonth | 12,97 | 2225,18 | 88,86 |
| cMonth | 4,45 | 2225,23 | 88,91 |
| cMonth + Clan | 4,68 | 2227,15 | 90,83 |
| sMonth + Clan | 13,14 | 2227,51 | 91,20 |
| sMonth * Clan | 13,30 | 2229,98 | 93,66 |
| sMonth + cYear * Clan | 30,72 | 2234,03 | 97,71 |
| qMonth | 1,17 | 2236,13 | 99,81 |
| qMonth + Clan | 1,37 | 2238,42 | 102,10 |
| qMonth * Clan | 2,03 | 2240,64 | 104,32 |
| Constant | 0 | 2242,30 | 105,98 |
| Clan | 0,20 | 2244,59 | 108,27 |
| cMonth * Clan | 6,91 | 2250,96 | 114,64 |
| sMonth * cYear * Clan | 42,01 | 2304,43 | 168,11 |

*Note:* The abbreviations *l* indicates a linear effect of the variable, *c* a categorical effect, *s* a non-linear effect, *q* a quadratic and *r* a random effect. The best equivalent models are in bold.

**Appendix S7: Annual patterns estimated from models that did not include a clan effect**

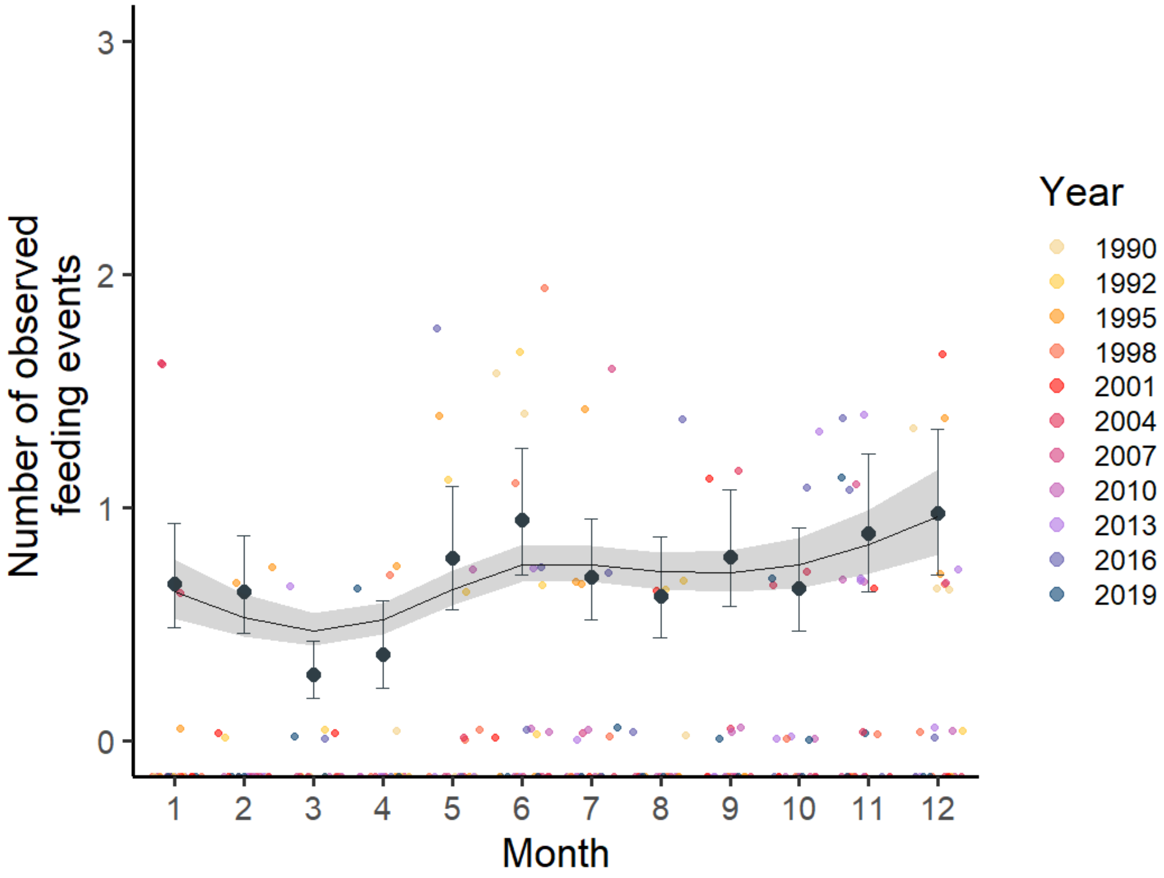


**Figure S1:** Annual pattern of the number of observed feeding events ($n_{feeding}$) estimated from a model with a categorical effect of month (*cMonth*; black dots with error bars) and from a model with a nonlinear effect of month (*sMonth*; black line and grey ribbon). Coloured dots represent raw data, that were jittered along the x and y-axis to improve visibility. The y-axis was cut at the 95% value of the data to facilitate visualization of estimated trends.

**Appendix S8: Variations in the annual patterns of the number of observed feeding events.**

Table S1: Values of the number of observed feeding events for different months over the years estimated from the *sMonth***sYear* model, and their 95% confidence intervals (between brackets).

| | **Model with interaction between *Month* and *Year*** | | | | |
|---|---|---|---|---|---|
| | **January** | **March** | **June** | **August** | **December** |
| **1990** | 1.25 | 0.85 | 1.17 | 1.06 | 1.18 |
| | [0.88, 1.79] | [0.65,1.12] | [0.97, 1.42] | [0.86, 1.31] | [0.81, 1.71] |
| **2004** | 0.62 | 0.47 | 0.74 | 0.74 | 0.99 |
| | [0.51, 0.76] | [0.40, 0.54] | [0.67, 0.82] | [0.66, 0.82] | [0.82, 1.19] |
| **2018** | 0.35 | 0.29 | 0.49 | 0.57 | 0.93 |
| | [0.23, 0.52] | [0.21, 0.41] | [0.42, 0.64] | [0.46, 0.70] | [0.65, 1.37] |

**Appendix S9: Estimates from model explaining the number of observed feeding events (*$n_{feeding}$*) by migratory prey presence (*$Prey_{max}$*).**

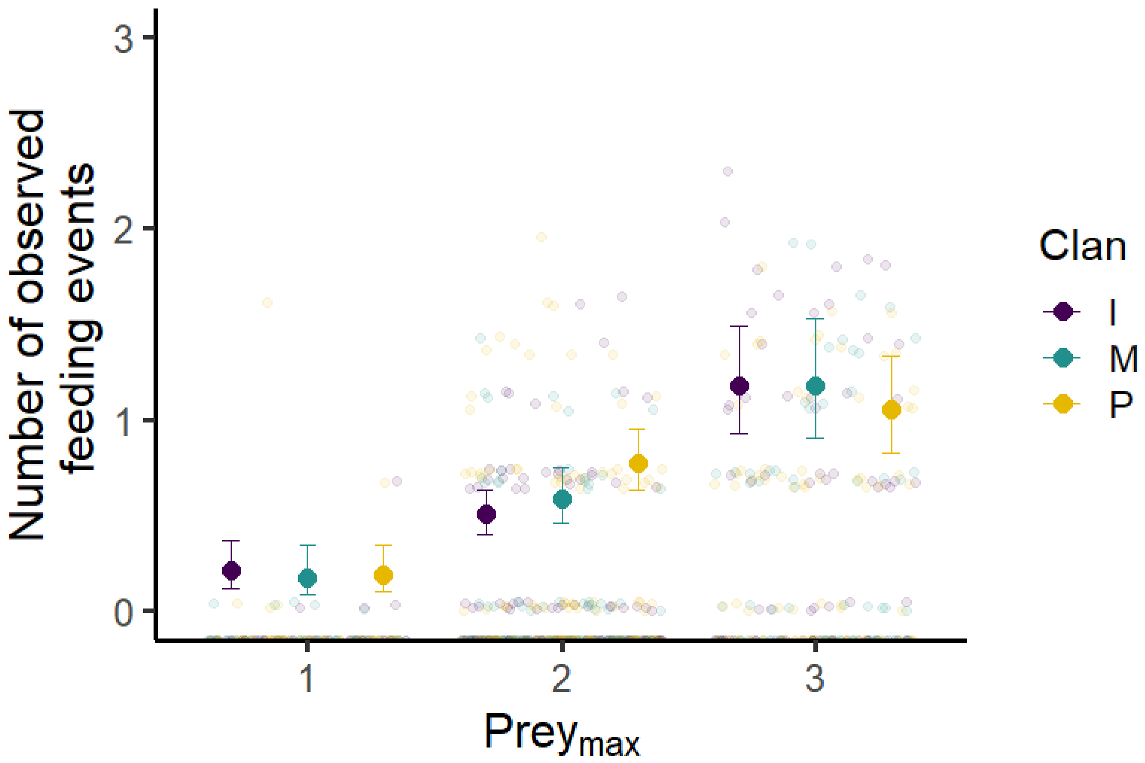


**Figure S1:** Number of observed feeding events per month as a function of $Prey_{max}$ (see Appendix S2, Table S1 for list of variables abbreviations and their meaning) for each clan. Estimates are indicated by large dots, along with their 95% confidence interval (error bars). The three clans showed similar trends. However, for Pool, $n_{feeding}$ is higher when $Prey_{max}$ is 2 and lower when $Prey_{max}$ is 3 compared to the other clans. Small coloured dots show the raw data, that were jittered along the x and y-axis to improve visibility. The y-axis was cut at the 95% value of the data to facilitate visualization of estimated trends.

**Appendix S10: Proportion of observed feeding events involving Thomson's gazelles over the years.**

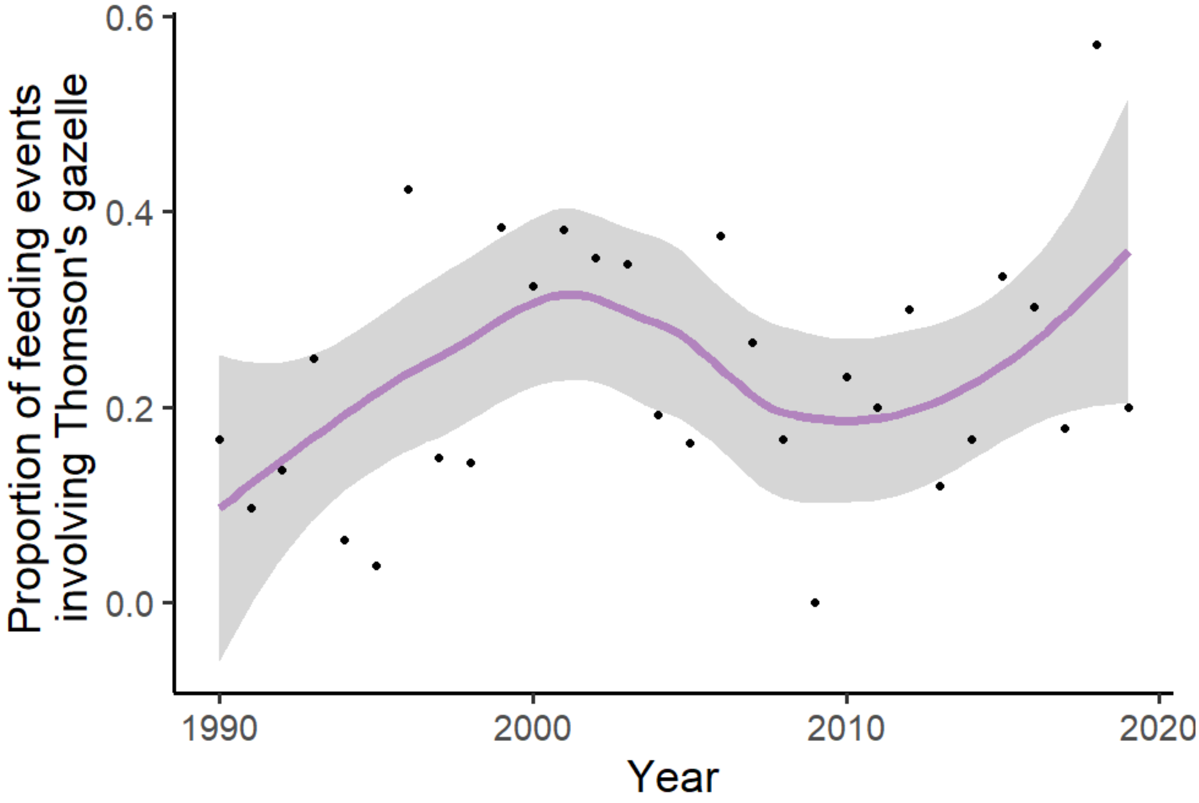


**Figure S1:** Proportion of observed feeding events involving Thomson's gazelles over the years. Black dots show the raw data, and the purple line and its ribbon the smooth of the trend The y-axis was cut at the 95% value of the data to facilitate visualization of estimated trends.
